# PI3K/AKT signaling allows for MAPK/ERK pathway independency mediating dedifferentiation-driven treatment resistance in melanoma

**DOI:** 10.1101/2022.07.08.499090

**Authors:** Eyleen Corrales, Ella Levit-Zerdoun, Patrick Metzger, Ralf Mertes, Ariane Lehmann, Julia Münch, Steffen Lemke, Silke Kowar, Melanie Boerries

**Affiliations:** Institute of Molecular Medicine and Cell Research (IMMZ), University of Freiburg, Stefan-Meier- Str. 17, 79104 Freiburg, Germany; Institute of Medical Bioinformatics and Systems Medicine (IBSM), Medical Center-University of Freiburg, Faculty of Medicine, University of Freiburg, Breisacherstr. 153, 79110 Freiburg, Germany; Faculty of Biology, University of Freiburg, Schänzlestr. 1, 79104 Freiburg,Germany; German Cancer Research Center (DKFZ), Im Neuenheimer Feld 280, 69120 Heidelberg, Germany; German Cancer Consortium (DKTK), Freiburg, Germany

**Keywords:** melanoma, MAPK/ERK, PI3K/AKT, dedifferentiation, quiescence, stemness, migration

## Abstract

Current therapeutic management of advanced melanoma patients largely depends on their BRAF mutation status. However, the vast heterogeneity of the tumors hampers the success of therapies targeting the MAPK/ERK pathway alone. Dissecting this heterogeneity will contribute to identifying key players in the oncogenic progression to tailor more effective therapies. We performed a comprehensive molecular and phenotypic characterization of a panel of patient-derived BRAF^*V600E*^-positive melanoma cell lines. Transcriptional profiling was used to identify groups of coregulated genes whose expression relates to an increased migratory potential and a higher resistance. A decrease in sensitivity to MAPK/ERK pathway inhibition with vemurafenib or trametinib corresponded with an increasing quiescence and migratory properties of the cells. This was accompanied by the loss of transcriptional signatures of melanocytic differentiation, and the gain of stem cell features that conferred highly-resistant/mesenchymal-like cells with increased xenobiotic efflux capacity. Nevertheless, targeting of the implicated ABC transporters did not improve the response to vemurafenib, indicating that incomplete BRAF inhibition due to reduced drug uptake is not a main driver of resistance. Rather, indifference to MAPK/ERK pathway inhibition arose from the activation of compensatory signaling cascades. The PI3K/AKT pathway in particular showed a higher activity in mesenchymal-like cells, conferring a lower dependency on MAPK/ERK signaling and supporting stem-like properties that could be reverted by dual PI3K/mTOR inhibition with dactolisib. In case of MAPK/ERK independency, therapeutic focus may be shifted to the PI3K/AKT pathway to overcome late-stage resistance in melanoma tumors that have acquired a mesenchymal phenotype.

## Background

Skin cancer is the most common malignancy, with melanoma representing only a 1-4% of all cutaneous neoplasia. Despite its relatively low incidence, malignant melanomas account for the majority of skin cancer related deaths. Although it has a good prognosis when diagnosed at early stages (with a 5-year survival rate of >90% at stage I), advanced stages of this neoplasia are very aggressive (with a 5-year survival rate of only 10-15% at stage IV), showing frequent metastasis and low treatment response (1). Available therapies for melanoma have considerably improved with the discovery of driver mutations in the B-rapidly accelerated fibrosarcoma (BRAF) gene. These mutations, harbored in >50% of melanomas, lead to the constitute activation of BRAF monomers (2,3). Consequently, hyperactivation of the mitogen-activated protein kinase/extracellular signal-regulated kinase (MAPK/ERK) pathway occurs, promoting proliferation, dedifferentiation, and survival of melanoma cells (4,5). Despite great progress has been achieved through the use of small-molecule inhibitors targeting mutated forms of BRAF and other MAPK/ERK pathway components, acquired resistance remains a main challenge.

The feasibility of developing “one-size-fits-all” therapies with durable responses is hindered by the large heterogeneity in melanoma. The disease has one of the highest frequencies of somatic mutations from all cancers (6), which results in a large intra- and inter-tumor variability (7). This underlies different cell states with divergent potentials to evolve into drug-tolerant or drug-resistant phenotypes. Hence, understanding the molecular basis beneath individual variation in oncogenic progression and treatment response is required to develop more tailored therapeutic approaches.

Here, in a systems biology approach, we aimed to dissect the behavioral and molecular heterogeneity in different patient-derived BRAF^*V600E*^-positive malignant melanoma cell lines (MaMel), with the aim to pinpoint key players implicated in the development of resistance to MAPK/ERK pathway inhibition. We infer a multistage and multifactorial nature of the oncogenic progression during the acquisition of drug resistance in melanoma, marked by the loss of cell differentiation, increased quiescence and migratory potential, reduced dependency on MAPK/ERK pathway, and activation of phosphatidylinositol 3-kinase/protein kinase B (PI3K/AKT) signaling. Our results raise considerations about the limited benefits of single agent therapies in advanced stages of the disease, and suggests that future efforts should be focused on the development of personalized combinatorial therapies.

## Methods

### Cell culture

Patient-derived BRAF^*V600E*^-positive malignant melanoma cell lines used in this study were provided by Prof. Dr. Stefan Eichmüller. The primary cultures were established at the German Cancer Research Center (DKFZ), Heidelberg, from skin (MaMel19, MaMel21, MaMel63a) or mediastinum (MaMel86b) metastases from female donors at stage IV of the disease (8).

The cell lines were maintained at 37 °C in a humidified atmosphere of 5% CO_2_ using RPMI-1640 medium supplemented with 10% FCS, 1% L-glutamine, and 1% penicillin-streptomycin. Pharmacological inhibition experiments were performed using 3 µM of vemurafenib (PLX4032), 5 nM of trametinib (GSK1120212), 3 µg/ml of dactolisib (BEZ235), or an analog volume of vehicle (0.3% of DMSO) as control, for the time indicated for each experiment.

### Transcriptome profiling and bioinformatic analyses

Total RNA was isolated using the Universal RNA Purification Kit (Roboklon), according to the manufacturer ‘s instructions. Microarrays-based gene expression profiling of MaMel cell lines was performed with Illumina HumanHT-12 v4 BeadChips, at the Genomics and Proteomics Core Facility of the DKFZ, Heidelberg. Each cell line was assessed in triplicates. Quantile normalization was performed using the Illumina BeadStudio software. A matrix with the normalized expression values for each cell line can be found in the Supplementary Table S1.

Gene expression patterns of MaMel cell lines were graphically depicted using a self-organizing maps (SOM) machine learning approach, as implemented in the som R package (9). This method allows dimensionality reduction, by assigning genes with similar expression patterns across the samples into clusters known as meta-genes. The meta-genes are subsequently arranged in a two-dimensional grid based on the similarity of the expression profiles (10). In the current study, SOM were constructed based on the normalized expression data from the cell lines using a two-dimensional grid of K = 40×40 meta-genes, for an average of n ≈ 12 genes per meta-gene with the Gaussian neighborhood function. Visualization was performed with a custom R code.

Groups of co-regulated genes across the different melanoma cell lines were identified through soft clustering analysis, using the Mfuzz R package (11). To this end, the fuzzy c-means algorithm was implemented in Euclidean space to cluster the top 30% of genes with the highest variance across the MaMel cell lines. The optimal number of clusters (c = 2) was defined by visual inspection for a plateau in the minimum centroid distance within a given range of cluster numbers (c = 2–10). A minimum membership score of 0.7 was required to assign each gene to a particular cluster.

Functional annotation of groups of genes was performed through over-representation analysis (ORA) with the R package clusterProfiler (12). Testing was performed for functional categories with a set size of 10-500 genes, and these were considered to be significantly over-represented with a q-value cutoff of 0.1. TRRUST v2 database (13) was used as reference to assess over-representation of candidate target genes of transcription factors. To test for enrichment of transcriptional signatures of interest in the different cell lines, gene set variation analysis (GSVA) was performed using the GSVA R package (14). Enrichment score (ES) values were calculated as the magnitude difference between the largest positive and negative random walk deviations.

To approach the degree of differentiation and the presence of unique malignant properties in the different cell lines, information properties of the transcriptomes (i.e., diversity and specialization) were estimated as previously described (15). An adaptation of Shannon ‘s entropy was implemented to determine the diversity of the transcriptomes (H), providing a minimum value from zero when only one gene is transcribed for a given sample, while reaching a maximum score when all genes are transcribed at the same frequency. The average specificity of the genes was used as an estimate of transcriptome specialization (δ), providing a minimum value of zero if all genes are expressed at the same frequency in all samples, while reaching a maximum score when genes are exclusively expressed in a particular sample.

### Flow cytometry

Proliferation rate of the cells was evaluated by flow cytometry measuring the retention of the Cell Proliferation Dye eFluor 670 (eBioscience), which binds to primary amines from the cells and retained in less proliferative cells. To this end, 1×10^5^ cells were stained for 10 min at 37 °C with 5 μM of the fluorescent dye. Unbound dye was then removed by washing the cells three times with supplemented cell culture medium. The cells were seeded, and allowed to attach for 6 h before inhibitor treatment. After the appropriate stimulation time, Annexin V and propidium iodide (PI) co-staining was performed to evaluate cell viability in parallel. To this end, cell culture supernatants containing all non-attached dead cells were first collected. The cells were then harvested, and pooled back together with their corresponding supernatants. The cells were pelleted and resuspended in an Annexin V-FITC dilution (1:25; BD Bioscience) on 1 × Binding Buffer. After 10 min incubation at room temperature, unbound Annexin V was washed with Binding Buffer, and the cells were resuspended in a PI dilution (1:50; BD Bioscience) and subjected to flow cytometry quantification.

Cell cycle phases were resolved by intracellular co-staining of Ki67 proliferation marker, phosphorylated histone H3, and DNA content with 4′,6-diamidino-2-phenylindole (DAPI). To this end, 2 ×10^5^ cells were harvested and permeabilized with 70% ice-cold ethanol, added drop-wise while vortexing the cell suspension. The samples were stored at −20 °C for at least 4 h, to ensure appropriate permeabilization. Blocking was performed for 30 min at room temperature, with a blocking solution of 3% FCS on PBS. The cells were then stained for 1 h at room temperature with PE-conjugated anti-Ki67 (1:100; SolA15; eBioscience), and Alexa Fluor 647-conjugated anti-phospho(Ser28)-histone H3 (1:100; HTA28; eBioscience) monoclonal antibodies, diluted in blocking solution. The cells were then rinsed with blocking solution, and resuspended in 250 µl of 1 µg/ml DAPI solution and subjected for flow cytometry analysis.

To study cell cycle kinetics, newly synthesized DNA was labeled using the Click-iT Plus 5-ethynyl-2’-deoxyuridine (EdU) Alexa Fluor 647 flow cytometry assay kit (ThermoFisher Scientific) according to manufacturer ‘s instructions, incubating for 1 h with EdU prior to staining. The cells were resuspended in 250 µl of 1 µg/ml DAPI solution and subjected for flow cytometry analysis.

Populations of stem cells were detected based on their dye efflux capacities using the side population (SP) assay. To this end, 1×10^5^ cells were harvested and stained in the dark with 10 µM Vybrant DyeCycle Violet (DCV; ThermoFisher) for 90 min at 37 °C, in the absence or presence of the ABC transporter inhibitors verapamil (200 µM; Sigma-Aldrich) and Ko143 (1 µM; Sigma-Aldrich). After staining, the cells were washed with ice-cold PBS, and resuspended in a 0.5 µg/ml solution of 7-amino-actinomycin D (7-AAD; eBioscience) to label and exclude dead cells from the analysis. The cells were kept on ice and analyzed with flow cytometry.

Flow cytometry acquisition from all experiments was performed on the BD LSR-II analyzer (BD Biosciences), using the DB FACSDiva software. A minimum of 10 000 events were acquired for every assessed condition. Data analysis was done using FlowJo 10.3 software, excluding debris and cell aggregates based on FSC/SSC characteristics.

### Real-time cell analysis (RTCA)

The xCELLigence System (ACEA Biosciences) was used to monitor cell migration (chemotaxis) and invasion of the cell lines in real-time. Experiments were performed using 16-well CIM plates, which consist of Boyden chambers with microelectrodes attached to the underside of the microporous membrane allowing an impedance-based detection of migrating/invading cells. For invasion experiments, the transwell inserts from the CIM plates were coated with 20 μl of growth-factor-reduced Matrigel (BD Biosciences), at a concentration of 250 μg/ml. Coated chambers were incubated for 4 h at 37 °C to allow homogenous polymerization of the Matrigel. The transwells were kept uncoated for migration experiments. Cells were resuspended in serum-free medium and seeded in the upper chambers, applying 2×10^4^ cells in each. Lower chambers were filled with medium supplemented with 10% FCS to serve as chemoattractant. Cells were allowed to migrate for 24 h, automatically monitoring the impedance (expressed as Cell Index; CI) every 30 min within the first 6 h, and every 60 min for the remaining time.

### Live cell tracking

MaMel cells were plated on 24-well dishes at a density of 8×10^3^ cells/cm^2^ and allowed to attach overnight. Movement of the cells was then recorded for 48 h using the JuLI Stage real-time cell history recorder system (NanoEnTek), by automatically capturing images of the wells every hour. The trajectories of 80 randomly selected cells from each cell line (measured from four independent experiments) were manually tracked using the ImageJ plugin MtrackJ (16).

### MTT assay

MTT assays were used to determine the concentration of inhibitors required to cause 50% growth inhibition (IC50) on MaMel cells. To this end, 1×10^4^ cells were plated in 96-well plates and allowed to attach overnight. Dilution series of the inhibitors were prepared and the cells were treated for 48 h, with every concentration performed in triplicates. The viability of the cells was then evaluated by incubating at 37 °C for 2.5 h with 200 mg/ml MTT (Calbiochem). The MTT solution was removed afterwards, and 200 µl of DMSO were added to every well. Absorbance was measured at 565 nm using an Infinite M200 PRO (Tecan) microplate reader. Measurements were normalized to the corresponding volume-matched DMSO controls, as maximal viability. Curves were fitted and IC50 calculated in Prism 8 (GraphPad), using a three-parameter nonlinear regression analysis.

### Western blot analysis

Cells were washed with PBS and then lysed by scrapping in 1 × RIPA buffer, supplemented with cOmplete™ EDTA-free protease inhibitor cocktail (Roche). Protein extracts were incubated for 40 min at 4 °C and spun down at 17000 × g for 10 min to remove cell residues. Total protein concentrations were determined with the DC Protein Assay (Bio-Rad), as indicated by the manufacturer. The extracts were then boiled for 5 min at 95 °C with 1 × Laemmli buffer in reducing conditions. Samples were stored at −80 °C until further use.

Protein lysates were resolved by loading 20 µg of total extracts in 12.5% SDS-PAGE, run at 150 V on 1 × TGS buffer (25 mM tris, 192 mM glycine, SDS 0.1%). Proteins were transferred onto methanol-activated Immobilon-P PVDF membranes (Merk Millipore) using a Mini-Transblot Cell (Bio-Rad), run for 2 h at 100 V and 4 °C on 1 × transfer buffer (25 mM tris, 200 mM glycine, 20% methanol). Membranes were blocked for 1 h at room temperature with 5% non-fat milk in 1 × TBS-T solution (20 mM tris, 150 mM NaCl, 0.1% Tween-20; pH 7.54).

Primary antibody dilutions were prepared in 1 × TBS-T containing 0.5% BSA and 0.1% NaN3. Membranes were probed overnight at 4 °C with antibodies against the following epitopes: phospho(Ser217/221)-MEK1/2 (1:1000; 41G9; Cell Signaling), MEK1/2 (1:1000; L38C12; Cell Signaling), phospho(Thr44/Tyr42)-ERK1/2 (1:1000; 9101; Cell Signaling), Erk1/2 (1:1000; L34F12; Cell Signaling), phospho(Ser473)-AKT (1:2000; D9E; Cell Signaling), AKT (1:2000; 40D4; Cell Signaling), phospho(Ser235/236)-S6 ribosomal protein (1:1000; 91B2; Cell Signaling), S6 ribosomal protein (1:1000; 54D2; Cell Signaling), BIM (1:1000; C34C5; Cell Signaling), FoxO1 (1:1000; C29H4; Cell Signaling), PTEN (1:1000; Cell Signaling). All membranes were probed with GAPDH (1:3333; 9484; Abcam) as loading control. After three washing steps with 1 × TBS-T, the membranes were incubated for 1 h at room temperature with the appropriate peroxidase-conjugated secondary antibodies, diluted in 1 × TBST with 5% non-fat milk: anti-rabbit IgG (1:10,000; NA934V; GE Healthcare) and anti-mouse IgG (1:10,000; NXA931V; GE Healthcare).

Chemiluminescence detection was done in the FUSION FX imaging system (Vilber Lourmat) using the SuperSignal West Pico Luminol/Enhancer solution (Thermo Scientific). The Fusion Capt Advance software (Vilber Lourmat) was used for densitometric analysis. The optical density of each band was normalized to the corresponding GAPDH loading control detected on the same membrane.

### Immunofluorescence staining

To evaluate the proliferative state of the cells, staining of the proliferation marker Ki67 was performed. To this end, 2.5 × 10^5^ cells were seeded in sterile cover slips placed at the bottom of 12-well dishes, and were allowed to attach overnight. Cells were then washed with PBS, fixed for 10 min with ROTI Histofix 4% solution (Carl-Roth), and permeabilized for 5 min with 0.2% Triton-X. After three washing steps with PBS, the slips were incubated for 1 h in the darkness with eFluor 450-conjugated anti-Ki67 (1:100; SolA15; eBioscience), and FITC-conjugated anti-α-tubulin (1:50; DM1A; Sigma) monoclonal antibodies. Washing steps were then repeated, and nuclear staining was then carried out for 15 min with a 0.5 µg/ml solution of 7-AAD (eBioscience). After three final washing steps, the cover slips were mounted using Dako Mounting Medium (Agilent Technologies). Acquired microscopy images are shown in pseudo-colors for a better visualization of Ki67 proliferation marker.

### Statistical analyses

Technical replicates within every experiment were averaged, and the average values from at least two independent experiments were used for statistical analysis. The exact number of independent experiments considered for every analysis is provided in the corresponding figure legends. Prism 8 software (GraphPad) was used to perform one- or two-way analysis of variance (ANOVA), followed by Tukey post-test to calculate statistical significance of the magnitude of changes between the experimental conditions. In all analyses, changes were considered significant at a p-value cutoff of 0.05. All data are presented as the mean ± SD of the experiments.

## Results

### Transcriptional profiles reflect the heterogeneity in the proliferative/invasive behaviors of BRAF^*V600E*^ mutant melanoma cell lines

To dissect the heterogeneity in malignant melanoma, and to contribute to the understanding of the molecular basis that could underlie different levels of resistance to MAPK/ERK pathway inhibition, we performed an in-depth comparison of four different patient-derived primary cell lines, all of them obtained from late-stage donors (stage IV) and positive for the BRAF^*V600E*^ mutation (8,17), albeit with marked morphological differences. Two of the cell lines (MaMel63a and MaMel86b) displayed a phenotype similar to that of normal melanocytes, with elongated dendrite-like protrusions, while the others (MaMel19 and MaMel21) exhibited a more dedifferentiated morphology (Fig. 1A).

**Fig. 1.**
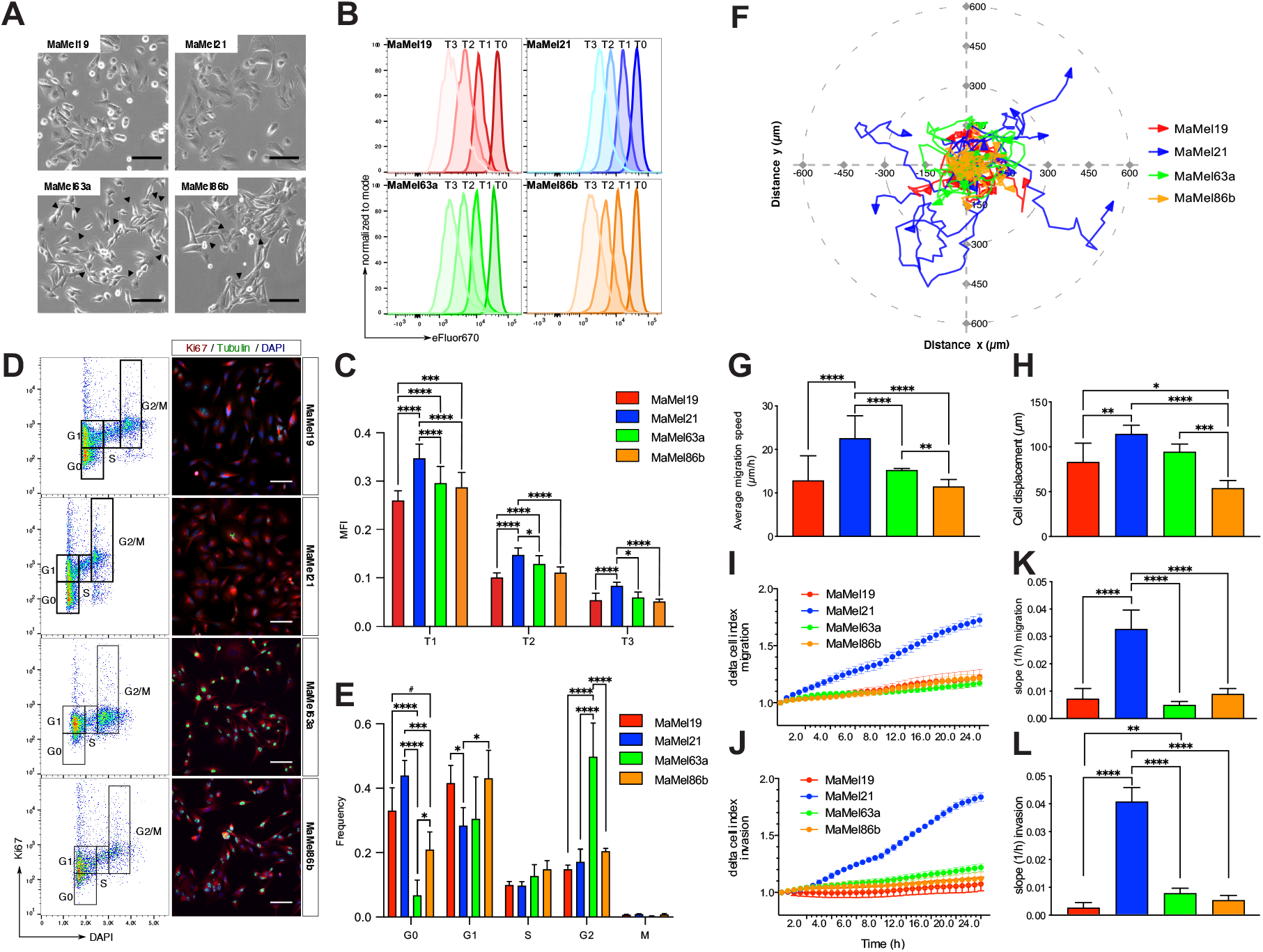
Motility and proliferation of BRAF^*V600E*^ mutant MaMel cells. **(A)** Phase contrast micrographs of MaMel cells. Arrows indicate dendrite-like protrusions in the cells. **(B)** Flow cytometric analysis of MaMel cell proliferation rate, detected as a reduction of the fluorescence intensity of eFluor 670 dye. Representative histograms at different time points (T) are shown for each cell line. **(C)** Quantification of the median fluorescence intensity (MFI) of the eFluor 670 dye. Data from three independent experiments are shown (n = 3). **(D)** Determination of proliferative state of the cells. Left panel: representative flow cytometric analysis of Ki67 and DAPI intracellular co-staining, defining the cell cycle phases as indicated. Right panel: representative immunofluorescence images of Ki67 staining. Nuclei (DAPI) and cytoskeleton (tubulin) are stained for visualization of the cellular structure. **(E)** Quantification of the frequencies of cells in each cycle phase shown in (D). M-phase was distinguished from the G2-phase by a histone H3 staining (not shown). Data from five independent experiments are shown (n = 5). **(F)** Live cell tracking of MaMel cells. Representative polar plot showing ten randomly selected individual tracks for each cell line. **(G)** Speed (µm/h) and **(H)** net displacement (µm) of cells during 48 h, averaged from 80 individual cell tracks. (**I-J**) Representative real-time cell analysis (RTCA) of the **(I)** migratory and **(J)** invasive behavior of MaMel cells. (**K-L**) Quantification of the slopes (1/h) from curves of cell **(K)** migration and **(L)** invasion over time obtained by RTCA. Data from two independent experiments are shown (n = 2). All bar plots show the mean ± SD of the experiments. *p < 0.05, **p < 0.01, ***p < 0.001, ****p < 0.0001. Scale bars: 200 µm.

We first compared the proliferative and cell cycling properties of the cell lines using flow cytometry-based proliferation assays, allowing for the dilution of the eFluor 670 proliferation dye to monitor the proliferative rate of the cells over time (Fig. 1B-C). The detection of the proliferation marker Ki67 (Fig. 1D-E) as well as the incorporation of the nucleotide analogue EdU served to resolve cell cycle stages (Fig. S1A-B). We found that MaMel21 was the least proliferative among the four cell lines (Fig. 1B-C), with highest number of cells in a quiescent state (G0) (Fig. 1D-E). The other cell lines were equally proliferative over time (Fig. 1B-C) but differed in their cell cycle state distribution, wherein MaMel19 had a larger quiescent G0 population than MaMel63a and MaMel86b (Fig. 1D-E), while the two latter cell lines had significantly higher S-Phase populations (Fig. S1A-B). In turn, MaMel63a, had the smallest G0 population, indicating that this cell line was the least quiescent (Fig. 1D-E). Analysis of migratory and invasive properties of the cell lines using live cell tracking and real-time cell analysis (RTCA) further revealed that MaMel21 was the most migratory and invasive among the cell lines (Fig. 1F-L), with higher average migration speeds (Fig. 1G) and larger cell displacements over time (Fig. 1H). These results indicate a phenotype switch in this cell line towards higher motility and invasiveness, and reduced proliferative capacity.

Microarray-based transcriptomic profiling of melanoma cells was then performed, for an unbiased identification of the key molecular differences that may underlie the observed phenotypic heterogeneity. Principal component analysis (PCA) revealed higher transcriptome similarities among the less-quiescent cell lines, MaMel63a and MaMel86b, while a larger heterogeneity was observed for MaMel19 and MaMel21 (Fig. 2A). To further compare the overall transcriptional profiles and identify the most variable sets of genes across the different cell lines, self-organizing maps (SOM) were constructed. The analysis revealed five main clusters of genes with distinctive expression patterns (Fig. 2B, Supplementary Table S2A-F). Consistent with the observation of a more migratory phenotype for MaMel21, this cell line displayed a high expression of genes encoding for extracellular matrix (ECM) components and remodelers such as different types of collagen and serpins, cell adhesion molecules including several integrin subunits, cytoskeleton and cell contraction molecules such as actin, components of transforming growth factor-beta (TGF-β) signaling, markers of epithelial-to-mesenchymal transition (EMT) like fibronectin 1 (FN1), and other genes linked to an increased migration and invasiveness such as AXL receptor tyrosine kinase (AXL) (Fig. 2B: spot A, Supplementary Table S2A, Supplementary Table S3A). Some of these genes also showed an intermediate expression level in MaMel19. In turn, this cell line was mainly characterized by a high expression of genes involved in antigen processing and presentation via MHC such as those encoding for several human leukocyte antigens (HLA), interferon signaling components, as well as an elevated expression of different matrix metalloproteinases (MMPs) (Fig. 2B: spot B, Supplementary Table S2B, Supplementary Table S3B).

**Fig. 2.**
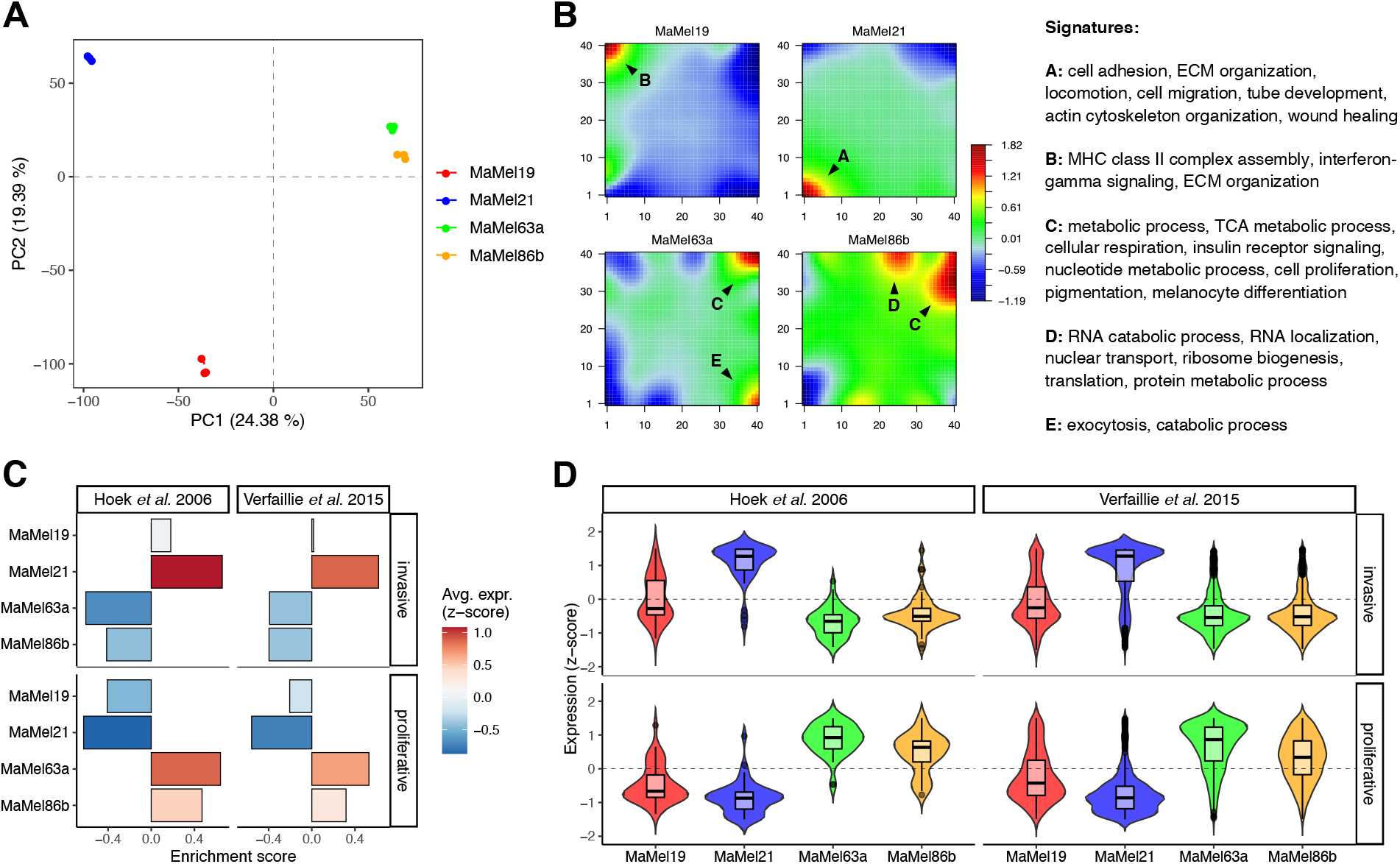
Transcriptional profiles in BRAF^*V600E*^ mutant MaMel cells. (**A**) Principal component analysis (PCA) for the transcriptomic profiles from MaMel cells. The projections of the first two principal components (PC1-PC2) are shown, with the percentage of variance explained by each one indicated in brackets. (**B**) Self-organizing maps of MaMel transcriptomes. Clusters of over- (red) and under- (blue) expressed genes are indicated with black arrows. Significantly enriched (p < 0.05) biological processes are annotated for each cluster. (**C**) Enrichment of proliferative and invasive melanoma signatures (21,22) in MaMel cells. (**D**) Normalized expression (z-score) from genes corresponding to the proliferative and invasive melanoma signatures.

On the other hand, MaMel63a and MaMel86b shared an elevated expression of genes implicated in different metabolic processes such as the tricarboxylic acid (TCA) cycle, nucleotide metabolism, cholesterol biosynthesis, insulin signaling pathway, and genes related to melanogenesis and pigmentation (Fig. 2B: spot C, Supplementary Table S2C-D, Supplementary Table S3C-D). In addition, MaMel86b also showed a strong up-regulation of genes related to RNA processing and ribonucleoprotein (RNP) complex biogenesis (Fig. 2B: spot D, Supplementary Table S2E, Supplementary Table S3E). Although no clear enrichment of proliferation-related processes was found that explained the lower quiescence of these two cell lines, their shared spot of up-regulated genes (Fig. 2B: spot C) included factors commonly associated with a more proliferative and less invasive phenotype, such as SRY-box transcription factor 10 (SOX10), melanocyte inducing transcription factor (MITF), and the MER proto-oncogene tyrosine kinase (MERTK) (18). Furthermore, negative regulators of the cell cycle progression, such as the cyclin-dependent kinase inhibitor p21 (CDKN1A) and the G0/G1 switch gene 2 (G0S2) (19,20), were found to be highly expressed in MaMel21 (Fig. 2B: spot A).

Gene Set Variation analysis (GSVA) was performed to compare the enrichment among MaMel cells of different transcriptional signatures previously reported for proliferative or invasive melanoma phenotypes (21,22). Consistent with our previous findings, a high enrichment of the invasive signatures in MaMel21 was found, with the vast majority of the genes showing increased expression in this cell line (Fig. 2C-D). In turn, MaMel19 had intermediate levels of enrichment of these signatures. By contrast, MaMel63a and MaMel86b exhibited a high expression of proliferative signature genes, with a slightly stronger enrichment in the former cell line. Altogether, these results evidence the heterogeneity in BRAF^*V600E*^ mutant melanoma cells, with transcriptional profiles that reflect the proliferative/migratory potentials of the tumors.

### Transcriptomic analysis suggests a multifactorial nature of resistance

Targeted therapy with small-molecule kinase inhibitors has revolutionized cancer treatment; however, their effects are frequently challenged by multifactorial resistance mechanisms, which may be of intrinsic and/or acquired nature. We therefore explored whether the observed phenotypic and transcriptional differences in the melanoma cells corresponded with different degrees of resistance to MAPK/ERK pathway inhibition. To this end, we treated the cells with vemurafenib, which specifically targets BRAF^*V600E*^, and trametinib, an inhibitor of MEK kinase activity. Western blot analysis revealed that both inhibitors induced a significant reduction in MEK and ERK phosphorylation in all four cell lines (Fig. 3A). However, this decrease was generally lower for the most migratory cell line, MaMel21 (Fig. 3B), suggesting differences in sensitivity of MaMel cell lines to MAPK/ERK targeting.

**Fig. 3.**
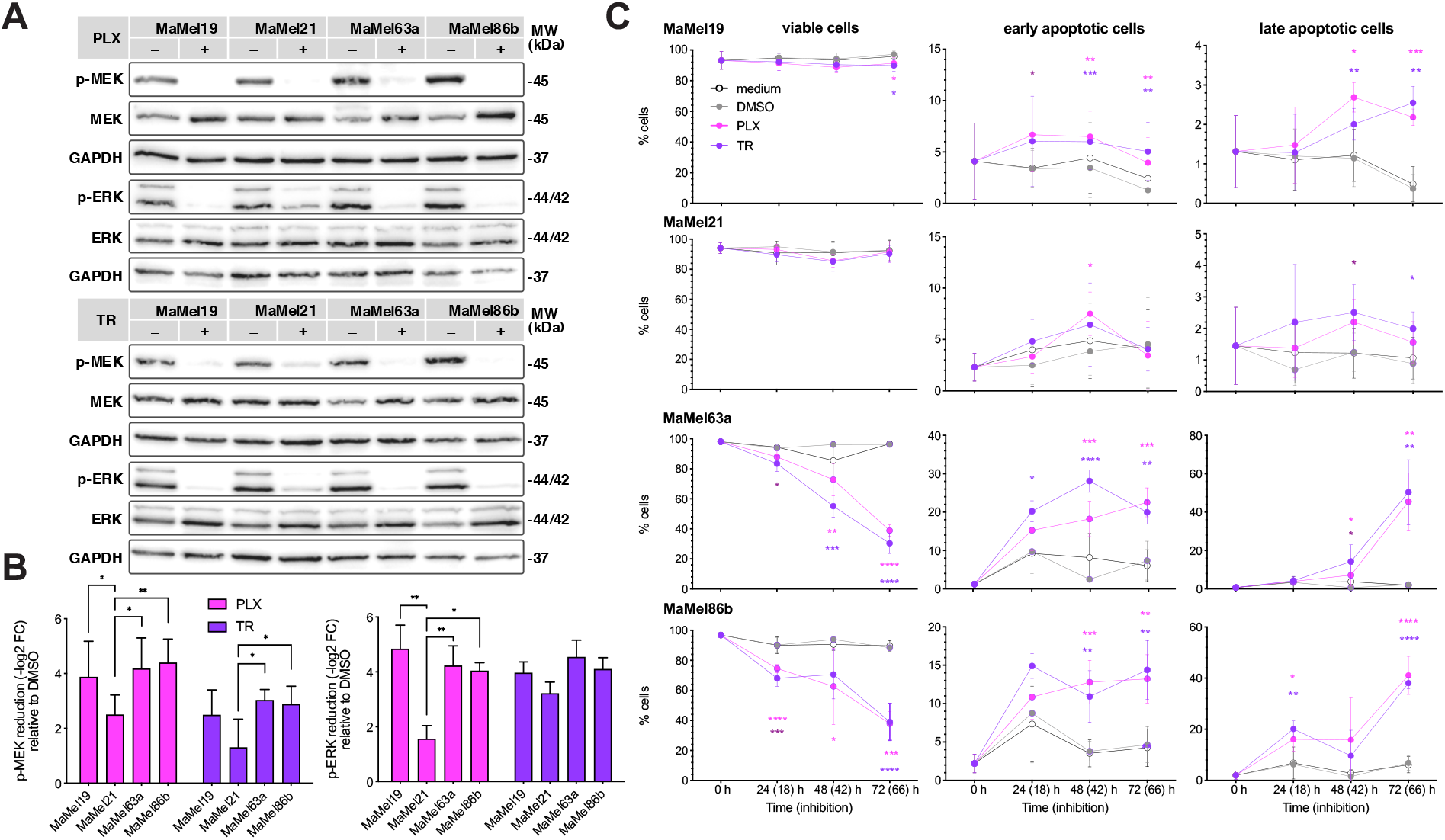
Response of BRAF^*V600E*^ mutant MaMel cells to MAPK/ERK pathway inhibition. **(A)** Representative Western blot analysis of MEK and ERK phosphorylation in MaMel cells treated with vemurafenib (3 µM, PLX; upper panel) or trametinib (5 nM, TR; lower panel). The phosphorylated (p-) and total proteins are shown; GAPDH served as loading controls. (**B**) Densitometric analysis of Western blots showing the fold change reduction (-log_2_ FC) of p-MEK and p-ERK after treatment with PLX or TR, in comparison to the corresponding vehicle (DMSO)-treated cells. Data from six independent experiments are shown (n = 6). **(C)** Quantification of the apoptotic response of MaMel cells under inhibition with PLX (3 µM) or TR (5 nM). Annexin V and PI co-staining, detected by flow cytometry, was used to define viable and early and late apoptotic cell populations at indicated time points. Data from two independent experiments are shown (n = 2). Time points outside the brackets indicate the total time of cells in culture; time points in brackets indicate the inhibition time. All bar and dot plots show the mean ± SD of the individual experiments. *p < 0.05, **p < 0.01, ***p < 0.001, ****p < 0.0001.

To address this in detail, we performed MTT-based viability assays, revealing a very high half-maximal inhibitory concentration (IC50) for vemurafenib in MaMel21 (mean IC50 = 58.96 µM) in comparison to the other cell lines, which ranged from 7 µM (MaMel19) to 0.5 µM (MaMel63a) (Fig. S1C-D). In the case of trametinib, although similar IC50 values were estimated for all cell lines (data not shown), it was noted that the viability of MaMel21 was barely impaired by the inhibitor, with around 80% of the cells still viable at the highest trametinib concentration (500 µM), whereas the viability of MaMel63a was greatly affected by high concentrations of the inhibitor (Fig. S1E-F). Annexin V-based apoptosis assays provided deeper insights on the different levels of sensitivity to inhibitor treatment, wherein MaMel21, but also MaMel19 were largely resistant to apoptosis under equimolar concentrations of either vemurafenib or trametinib even at later time points of inhibition. By contrast, MaMel63a and MaMel86b were highly prone to apoptosis (Fig. 3C, Fig. S2A), possibly due to elevated basal levels of BIM detected in these cells (Fig. S2B). We additionally performed flow cytometry-based proliferation assays, which revealed that proliferation upon vemurafenib and trametinib treatment was significantly inhibited over time in MaMel19 and MaMel21 (at 48 and 72 h) and to some extent in MaMel63a (at 72 h), but not in MaMel86b (Fig. S2C-D); however, these results should be interpreted with caution due to high apoptosis ratios in the two latter cell lines (Fig. 3C). Taken together, these results show that MaMel cell lines are highly heterogeneous in their response to targeted inhibition, wherein the most migratory cell line, MaMel21, is also highly resistant to BRAF and MEK inhibition. In turn, MaMel19 shows drug tolerance, as evidenced by a low apoptosis ratio in these cells, whereas the least quiescent cell lines, MaMel63a and MaMel86b, were treatment-sensitive.

It was noted that this variability in the response to therapeutic agents targeting the MAPK/ERK pathway is likely reflected at the transcriptional level. In particular, the first principal component of the PCA (PC1), which accounts for almost a quarter of the variance in gene expression, mirrored the different levels of sensitivity to MAPK/ERK pathway inhibition: MaMel21 was the most distinct cell line, and MaMel19 displayed an intermediate profile (Fig. 2A). This suggested the existence of prominent transcriptional differences underlying the degree of resistance of the tumor cells.

To better identify groups of co-regulated genes along the degree of resistance, we performed soft clustering for the top 30% (n = 5899) of genes with the highest variance across the cell lines. One cluster of genes showed a higher expression with increasing resistance levels to MAPK/ERK pathway inhibition (Fig. 4A: cluster 1). This cluster included genes encoding for proteins involved in vasculogenesis such as vascular endothelial growth factors (VEGFs) and platelet-derived growth factors (PDGFs) (23), cell migration like N-cadherin (CDH2), and stem cell population maintenance including fibroblast growth factor 2 (FGF2) and high mobility group box 2 (HMGB2) (24,25), likely reflecting an increasing metastatic potential and the gain of stem cell features (Supplementary Table S4A, Supplementary Table S5A). Likewise, hallmarks of p53 activity and immune-like response were also observed (Supplementary Table S6A). Concurrently, there was a reduction in the expression of genes implicated in melanocytic differentiation and oxidative phosphorylation (Fig. 4A: cluster 2, Supplementary Table S5B, Supplementary Table S6B). Well-known melanocytic genes, like melanocortin 1 receptor (MC1R), tyrosinase (TYR), dopachrome tautomerase (DCT), premelanosome protein (PMEL), and melan-A (MLANA), as well as their common upstream regulator MITF, were part of this cluster of genes (Supplementary Table S4B). This observation was consistent with an overall lower expression in the most resistant MaMel cell lines of MITF/SOX10-regulated signature genes (Fig. 4B), previously reported to be down-regulated in melanoma cohorts with higher metastatic potentials (21).

**Fig. 4.**
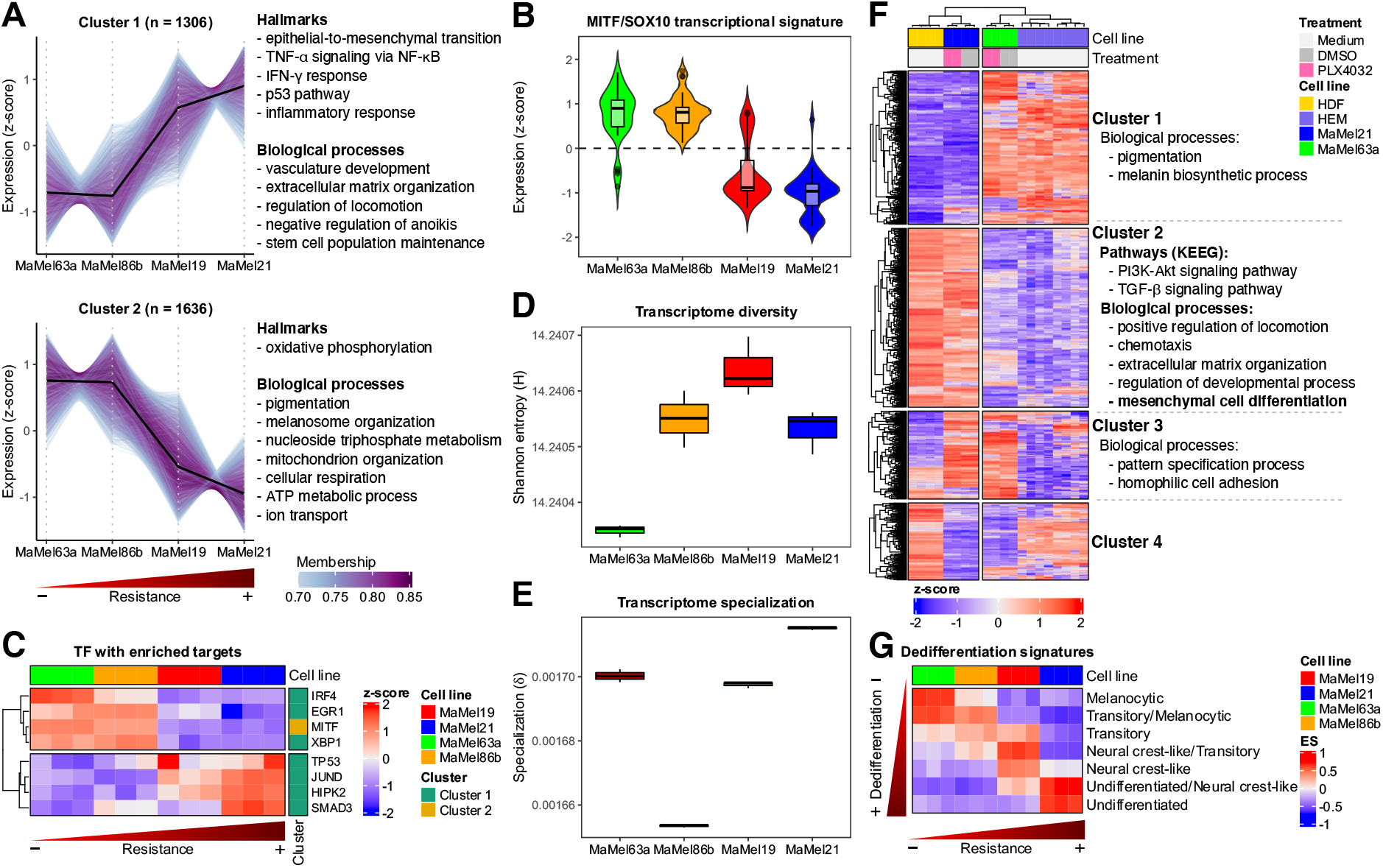
Identification of transcriptional signatures of resistance and differentiation in BRAF^*V600E*^ mutant MaMel cells. **(A)** Soft clustering based on the top 30% of genes with the highest variance across MaMel cell lines. The number of genes within every cluster is indicated in brackets; only genes with a minimum membership score of 0.7 are included. Some of the main significantly over-represented (q < 0.05) hallmarks and biological processes for every cluster are annotated next to the respective plot. **(B)** Normalized expression (z-score) from genes (n = 1549) corresponding to the MITF/SOX10 transcriptional signature (21). **(C)** Normalized expression (z-score) from transcription factors with enriched targets within the clusters identified on (A). **(D)** Transcriptome diversity, and **(E)** specialization of MaMel cells. **(F)** Heatmap displaying the normalized expression (z-score) of the 10% of genes with the highest variance across MaMel cell lines, human dermal fibroblasts (HDF), and human epidermal melanocytes (HEM). Some of the main significantly over-represented (q < 0.05) biological processes and pathways are annotated alongside the clusters of genes (identified by k-means clustering) with similar expression patterns across the samples. **(G)** Enrichment score (ES) of transcriptional signatures of progressive melanoma dedifferentiation.

To pinpoint master regulators that could orchestrate a transcriptional remodeling during acquisition of resistance, we checked for over-representation of target genes of transcription factors within the aforementioned clusters. The analysis provided further evidence that genes whose expression was reduced with increasing degree of resistance (Fig. 4C: cluster 2) mainly corresponded to targets of MITF, which in turn had a higher expression on the most sensitive cell lines (Fig. 4C, Fig. S3A). On the other hand, targets from several transcription factors were significantly enriched in the cluster of genes whose expression increased with resistance (Fig. 4C: cluster 1, Fig. S3A). This included p53 and its coactivator homeodomain-interacting protein kinase 2 (HIPK2), which had a higher expression in the most resistant cell lines and are likely involved in the transition to a slow-cycling phenotype (26). Moreover, the analysis also revealed an enrichment of targets of JunD, which has been suggested to protect from p53-mediated senescence and apoptosis (27), as well as targets of mothers against decapentaplegic homolog 3 (SMAD3) as a key transducer of TGF-β signaling. Other transcription factors showed an inverse expression pattern with their predicted target genes, suggesting a negative regulation. This included the interferon regulatory factor 4 (IRF4), a transcription factor known to be induced by SOX10 that acts as a negative regulator of IRF1 to repress melanoma immunogenicity (28). The expression of these transcription factors was further addressed in the publicly available dataset of Tsoi et al. (29), comprising 53 melanoma cell lines with different NRAS/BRAF mutation status and degrees of differentiation. This revealed expression patterns consistent with the degree of differentiation, with MITF and IRF4 showing higher expression levels in the more melanocytic-like cell lines (i.e., melanocytic and transitory subtypes). Simultaneously, JUND, HIPK3, SMAD3, and TP35 were generally highly expressed in the less differentiated states (i.e., neural crest-like and undifferentiated subtypes) (Fig. S3B).

As our analyses implied that the increasing resistance of the cells was accompanied by a reduction of their melanocytic features and the acquisition of migratory properties, the diversity and specialization of the transcriptomes from the different cell lines was estimated to approach their degrees of differentiation and gain of unique malignant features. Transcriptome diversity was quantified by an adaptation of Shannon ‘s entropy, as previously described (15), providing a value of zero when only one gene is transcribed for a given sample and a maximum score when all genes are transcribed at the same frequency. This analysis exposed MaMel63a as the cell line with the least diverse transcriptome, a particularity observed in more differentiated cells that usually require more limited transcriptional repertoires to fulfill their specific functions (30). As opposed, MaMel19 had the most diverse transcriptome (Fig. 4D). On the other hand, transcriptome specialization was estimated as the average gene specificity, whose value would be zero if genes were transcribed at the same frequency in all samples, while a maximum value would be reached if genes were exclusively expressed in a particular sample (15). The highest transcriptome specialization was observed for MaMel21, indicating a larger expression of more specific genes and suggesting the acquisition of additional properties that are not shared with the other cell lines (Fig. 4E).

To further investigate the degree of differentiation of the cells, and to determine whether they retained more melanocytic features or if they had acquired mesenchymal properties, previously generated RNA-seq profiles from the most diverging cell lines, MaMel63a and MaMel21, were compared to those from publicly available datasets from human epidermal melanocytes (HEM) (31,32), and human dermal fibroblasts (HDF) as prototypical mesenchymal cells (33). Hierarchical clustering based on the top 10% (n = 1549) of the genes with the highest variance across the different cell types confirmed a higher similarity of MaMel63a with HEM. Consistent with our previous observations, these cell lines shared a high expression of genes involved in melanin biosynthesis and pigmentation (Fig. 4F: cluster 1, Supplementary Table S7A, Supplementary Table S8A). On the contrary, MaMel21 showed greater similarities with HDF. A large set of genes involved in cell migration, extracellular matrix organization, and other processes shared a high expression in these cell lines (Fig. 4F: cluster 2, Supplementary Table S7B, Supplementary Table S8B). KEGG pathway analysis further suggested an up-regulation of PI3K/AKT and TGF-β signaling pathways in MaMel21 and HDF (Supplementary Table S9B). A smaller set of genes involved in development and chromatin organization, including several members of the homeobox family as well as histone encoding genes, shared an enhanced level of expression in these two melanoma cell lines (Fig. 4F: cluster 3, Supplementary Table S7C, Supplementary Table S8C). However, it should be noted that, according to the hierarchical clustering, the similarities between the tumor cell lines were largely outweighed by their respective similarity with HEM or HDF, even under BRAF inhibition with vemurafenib. GSVA confirmed a correspondence in the enrichment of signatures of progressive melanoma dedifferentiation reported by Tsoi et al. (29) with the increasing degree of resistance in the MaMel cells (Fig. 4G).

Taken together, these results imply that the acquisition of resistance in melanoma cells might be explained by the loss of differentiation and gain of cell pluripotency, followed by a trans-differentiation into a more mesenchymal-like phenotype, with the acquisition of migratory properties and the activation of alternative signaling pathways.

### Resistance to BRAF inhibition does not rely on efflux capacities

Based on our results showing a reduced differentiation and higher quiescence upon increasing resistance, and given that these properties are frequently endowed in cancer stem cells (34), we further explored the degree of stemness on the different MaMel cell lines. To this end, the side population (SP) assay was employed as a way to detect subpopulations of stem cells, based on their dye efflux capacities via ATP-binding-cassette (ABC) family transporter proteins (35). This revealed that MaMel21, unlike the other cell lines, has a considerably large SP comprising a subset of about half of the cells (Fig. 5A-B).

**Fig. 5.**
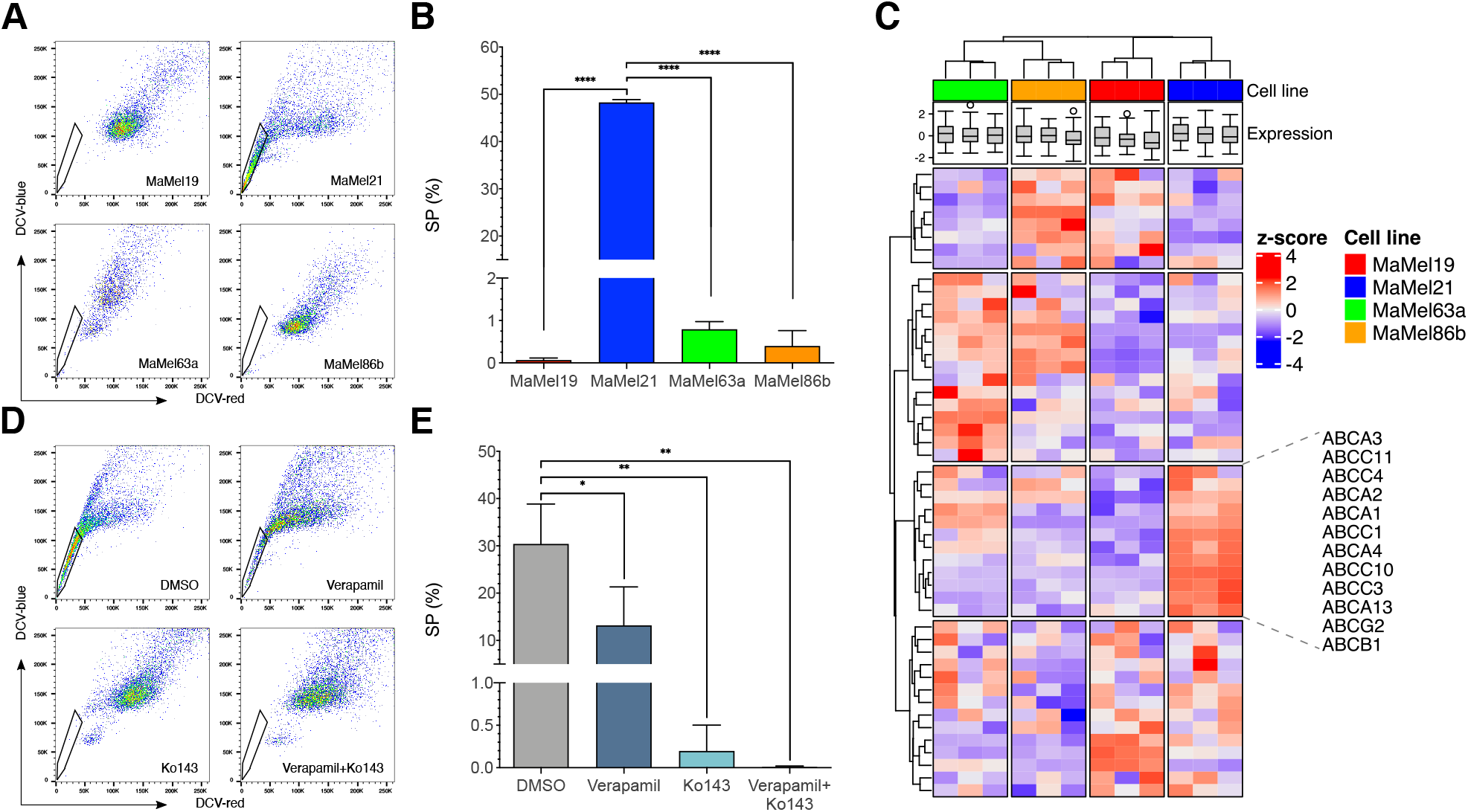
Efflux capacity of BRAF^*V600E*^ mutant MaMel cells. **(A)** Representative scatter plots of the side population (SP) assay detecting the efflux of DyeCycle Violet stain (DCV) in MaMel cells. The polygonal gates indicate the SP subsets (DCV-) for each cell line. **(B)** Quantification of the percentage of SP cells. **(C)** Heatmap displaying the normalized expression (z-score) of genes encoding for ABC transporter family members. **(D)** Representative scatter plots of the SP assay for MaMel21 after inhibition of ABC transporters with verapamil, Ko143, or the combination, in comparison to DMSO control. The polygonal gates indicate the SP subsets for each condition. **(E)** Quantification of the percentage of SP cells. All bar plots show the mean ± SD of three independent experiments (n = 3). *p < 0.05, **p < 0.01, ****p < 0.0001.

At the transcriptional level, we found that although the four cell lines did not differ in the overall expression of 49 genes encoding for ABC family members, a particular subset of transporters implicated in multidrug resistance was over-expressed in MaMel21 (Fig. 5C). This included ABCB1 and ABCG2, for which vemurafenib is considered to be a substrate (36). By treating MaMel21 with the ABC transporter inhibitors verapamil (which blocks several ABC drug transporters, with a main affinity for ABCB1), Ko143 (which selectively blocks ABCG2), or the combination, we explored their relevance in the efflux capacity of MaMel21. Although verapamil did not fully preclude the dye efflux from this cell line, Ko143 alone was sufficient to eradicate the SP, supporting a relevant role of the ABCG2 transporter in xenobiotic exclusion ability (Fig. 5D-E).

Given that MEK and ERK phosphorylation levels after vemurafenib treatment were slightly less diminished in MaMel21 than in the other cell lines (Fig. 3A), we investigated whether the greater resistance in this cell line could be explained by incomplete MAPK pathway inhibition due to a higher efflux capacity of the inhibitor, and whether ABC transporter inhibition could help to overcome resistance by enhancing the drug uptake. To this end, we performed time-course experiments of combined treatment with Ko143 and vemurafenib. However, ABCG2 inhibition with Ko143 did not sensitize MaMel21 or any other cell line to vemurafenib, inducing no further impact on cell proliferation (Fig. S4A-B) or viability (Fig. S4C-D).

In summary, these results indicate that, although resistant melanoma cells can bear stem-like properties with increased xenobiotic efflux capacities, targeting of ABC transporters has a limited potential to improve the response to BRAF targeted therapies.

### PI3K/AKT pathway activation underlies increased resistance to BRAF inhibition

As targeting of ABC transporters to prevent vemurafenib efflux did not improve the response of MaMel21 to this BRAF inhibitor, we speculated that this could reflect a lower dependency of this cell line on signaling through the MAPK/ERK pathway. We therefore checked the expression of genes from the MEK-dependent transcriptional signature reported by Pratilas et al. (37) as a readout of the transcriptional activity of the pathway (Fig. 6A). This evidenced an overall lower expression of these signature genes in MaMel21, as compared to the other cell lines, indicating a lower reliance of this cell line on the activity of the MAPK/ERK pathway. The activation state of the PI3K/AKT pathway was therefore explored as an alternative signaling cascade, motivated by our analysis indicating that this could be a feature shared among cells with mesenchymal properties (Fig. 4F).

**Fig. 6.**
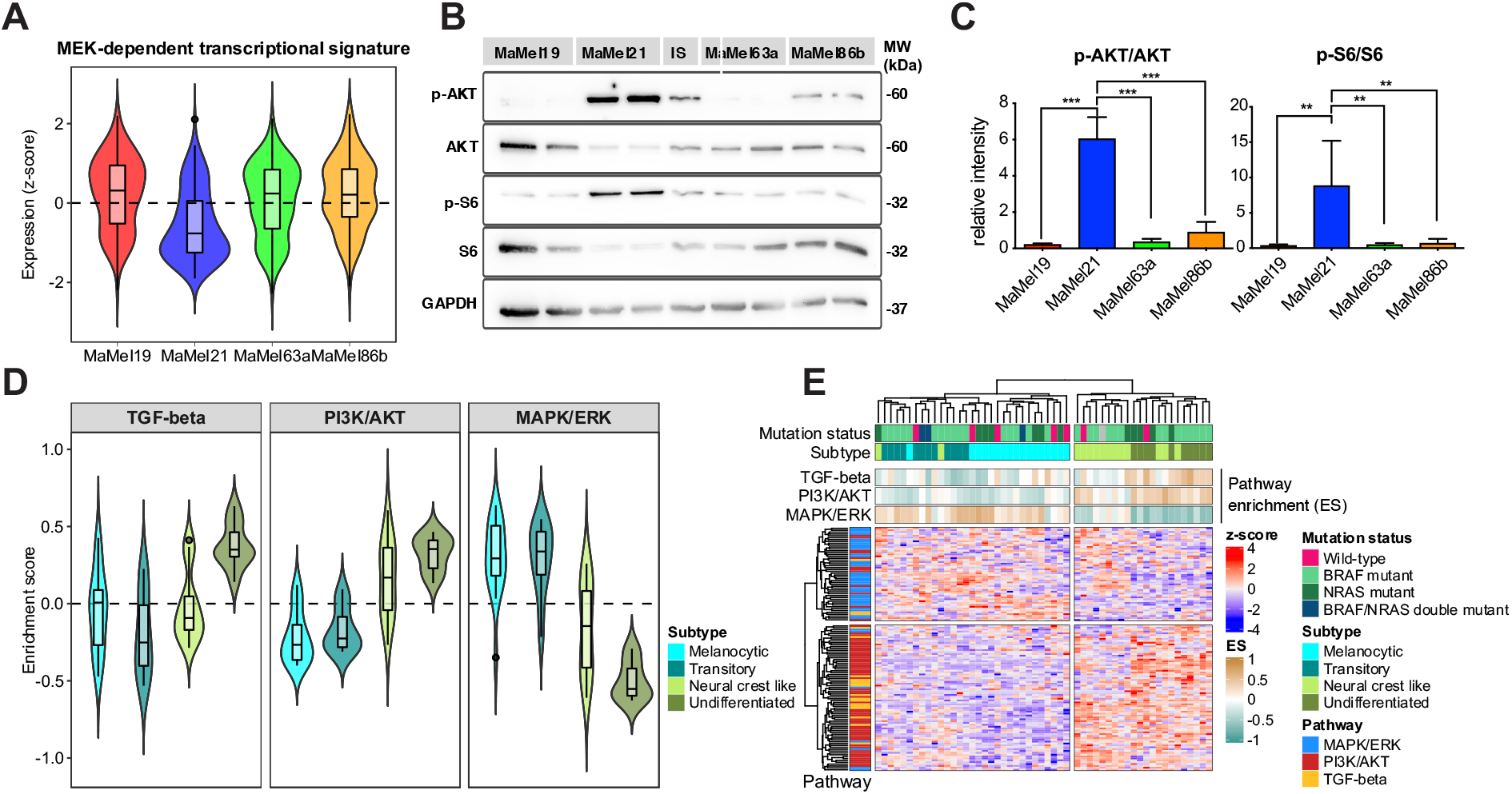
PI3K/AKT pathway activity in melanoma cell lines. **(A)** Normalized expression (z-score) from genes (n = 49) corresponding to the MEK-dependent transcriptional signature (37) in MaMel cell lines. **(B)** Representative Western blot analysis of AKT and S6 phosphorylation in MaMel cell lines. The phosphorylated (p-) and total proteins are shown; GAPDH served as loading control. **(C)** Densitometric analysis of Western blots showing the relative intensity from p-AKT and p-S6 to total AKT and S6, respectively. Data from three independent experiments are shown (n = 3). **(D)** Heatmap displaying the scaled expression (z-score) of signature genes from MAPK/ERK, PI3K/AKT, and TGF-β pathways in 53 melanoma cell lines from Tsoi et al. (29), with different NRAS/BRAF mutation status and degrees of differentiation The top annotations show the level of enrichment (ES) of the signatures in every cell line. **(E)** Enrichment of signatures from the MAPK/ERK, PI3K/AKT, and TGF-β pathways for the distinct differentiation subtypes within the aforementioned panel of melanoma cell lines.

By Western blot analysis, we detected a hyperphosphorylation of AKT in MaMel21, indicative of a constitutive activation of the pathway (Fig. 6B-C). In line with this, MaMel21 cells displayed the highest activity of the mammalian target of rapamycin (mTOR) pathway, a downstream target of phosphorylated AKT, indicated by higher phosphorylation of the ribosomal protein S6, a surrogate marker of mTOR pathway activation (Fig. 6B-C). Similarly, the levels of the downstream negatively regulated target forkhead box protein O1 (FoxO1; Fig. S5) were decreased in this cell line. AKT hyperphosphorylation in MaMel21 was likely explained by lack of phosphatase and tensin homolog (PTEN; Fig. S5), a negative regulator of PI3K signaling with a homozygous deletion reported for this cell line (17).

To determine if a reduced MAPK/ERK signaling and an increased activity of PI3K/AKT and TGF-β pathways were consistently observed upon loss of differentiation in other melanoma cell lines (Fig. 4F), we first derived signatures of transcriptional dependency on the pathways. The MAPK/ERK pathway-dependency signature was composed by signaling transducers and target genes under-expressed in the less-differentiated/highly-resistant melanoma cell line, MaMel21, in comparison to the more-melanocytic/drug-sensitive cell lines, MaMel63a and MaMel86b. In the case of PI3K/AKT and TGF-β signatures, these were composed by pathway components over-expressed in MaMel21, as compared to MaMel63a and MaMel86b. The enrichment of these signatures was addressed in the dataset of Tsoi et al. (29), revealing that cell lines corresponding to the melanocytic and transitory subtypes had higher levels of MAPK/ERK transcriptional activity, but a lower expression of signature genes from the PI3K/AKT pathway (Fig. 6D-E). This relation was progressively inverted with increasing loss of differentiation, shifting to an increased PI3K/AKT transcriptional activity and reduced MAPK/ERK signature expression (Fig. 6D-E). These changes were particularly evident in the undifferentiated state, which additionally showed high TGF-β pathway activity (Fig. 6E).

To further investigate the dependency of MaMel cells on the activity of MAPK/ERK and PI3K/AKT pathways, we assessed the changes in AKT and S6 phosphorylation after treatment with either the MAPK/ERK inhibitors vemurafenib or trametinib, or with the dual PI3K/mTOR inhibitor dactolisib. Western blot analysis revealed that levels of phospho-AKT were reduced in all cell lines under dactolisib treatment (Fig. 7A-B). Furthermore, they were equally sensitive to dactolisib (Fig. S1G-H). Notably, however, the levels of phosphorylated S6 in MaMel21 were largely affected by PI3K/mTOR inhibition and less so by MAPK/ERK inhibition (Fig. 7A-B). As phosphorylation of ribosomal protein S6 is considered a point of convergence from both signaling pathways, this indicates a higher reliance of this cell line on the PI3K/mTOR pathway activity. Moreover, the proliferation of MaMel21 was significantly affected by dactolisib after 66 hours of treatment (Fig. 7C, Fig. S2C-D). Remarkably, dactolisib almost completely eradicated the SP in MaMel21 after 66 h of treatment, suggesting the potential of PI3K/mTOR pathway inhibition to restrain some of the stem cell properties from the cell line (Fig. 7D-E).

**Fig. 7.**
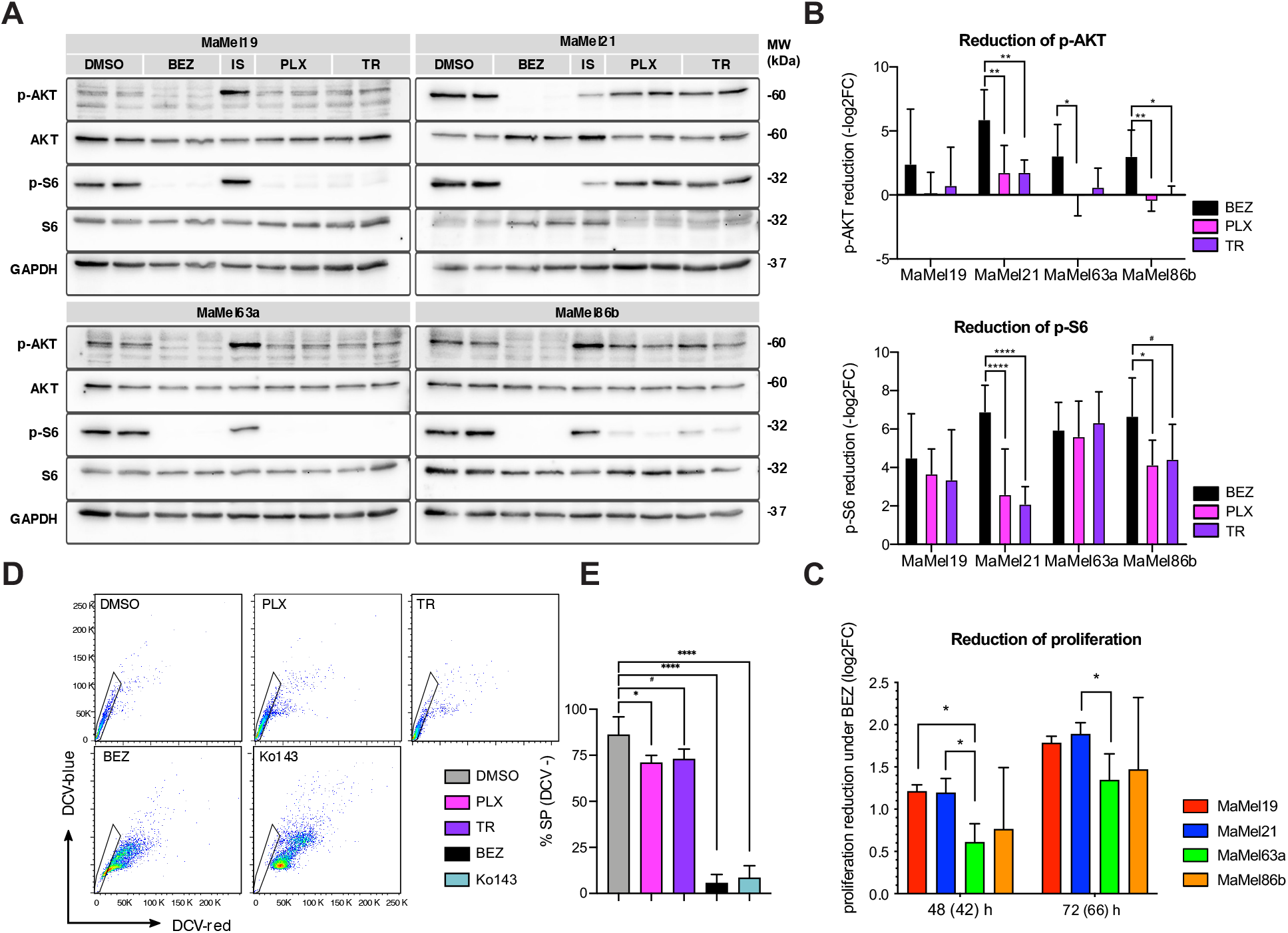
Response of BRAF^*V600E*^ mutant MaMel cells to PI3K/AKT pathway inhibition. **(A)** Representative Western blot analysis of AKT and S6 phosphorylation in MaMel cells treated with dactolisib (BEZ), vemurafenib (PLX), or trametinib (TR). The phosphorylated (p-) and total proteins are shown; GAPDH served as loading control. **(B)** Densitometric analysis of Western blots showing the fold change reduction (- log_2_ FC) of p-MEK and p-ERK after treatment with PLX or TR, in comparison to the corresponding vehicle treated cells. Data from five independent experiments are shown (n = 5). **(C)** Fold change reduction of the cell proliferation rate (log_2_ FC) of MaMel cells under inhibition with BEZ at the indicated time points. Data from three independent experiments are shown (n = 3). The time points outside the brackets indicate the total time of cells in culture. Time points in brackets indicate the inhibition time. **(D)** Representative side population (SP) assay detecting the efflux of DyeCycle Violet stain (DCV) in MaMel21 under inhibition for 66 h with PLX, TR, BEZ, or the ABCG2 inhibitor Ko143. The polygonal gates indicate the SP subsets (DCV-) in each condition. **(E)** Quantification of the percentage of SP in MaMel21 under inhibition. Data from five independent experiments are shown (n = 5). All bar and dot plots show the mean ± SD of the individual experiments. *p < 0.05, **p < 0.01, ***p < 0.001, ****p < 0.0001.

Altogether, these data suggest that loss of differentiation and acquisition of resistance to MAPK/ERK pathway inhibition is delineated by a reduction in the dependency on this pathway, with a concurrent increase in PI3K/AKT signaling that confers melanoma cells with stem-like properties.

## Discussion

Mutations in BRAF, particularly the V600E, are frequently harbored in melanoma patients (2). These genetic alterations are considered as one of the main drivers of the malignant transformation of melanocytes (38). Currently, the first-line treatment for advanced melanoma greatly depends on the mutation status of BRAF. In the presence of BRAF mutations, targeted therapies with BRAF inhibitors are usually administered to the patients, either as single-agents or in combination with MEK inhibitors. However, although this approach has considerably improved the prognosis for patients with metastatic melanoma, most of them relapse during treatment due to acquired resistance.

Here we show that, beneath the same BRAF mutation state, a large phenotypic and molecular heterogeneity can be found among late-stage malignant melanomas, which hinders the success of therapies solely targeting the MAPK/ERK pathway. This heterogeneity might explain why previous studies have disagreed on whether gene expression profiles correlate with the presence of activating mutations in BRAF or NRAS. While some investigations have described transcriptional signatures characteristic for mutated BRAF or NRAS tumors (39,40), larger cohorts have failed to identify any clear correlation between the transcriptional profiles and the mutational status of the cells (21). This lack of consistency might be reflecting the progressive acquisition of new malignant features and the activation of compensatory signaling cascades, allowing the cells to diminish their dependency on the MAPK/ERK pathway during late stages of malignant transformation, as shown here for a panel of morphologically-distinct patient-derived melanoma cell lines harboring the BRAF^*V600E*^ mutation.

We recognized in our panel of cell lines three previously reported main melanoma states, corresponding to melanocytic, intermediate, and mesenchymal-like phenotypes (41). The melanocytic-like state is characterized by a highly proliferative phenotype, with a high expression of genes involved in pigmentation and melanocytic differentiation. As opposed, the mesenchymal-like state reflects the loss of expression of melanocytic genes and an increased cellular quiescence. This is accompanied by the gain of migratory and vasculogenic properties, indicative of a higher metastatic potential of these cells. An intermediate state is characterized by the expression of immune-related transcriptional programs, and likely reflects the transition from the differentiated/proliferative state into an undifferentiated/migratory phenotype, sharing mixed features from the melanocytic- and mesenchymal-like subtypes. Hence, our observations support a correspondence of the transcriptional profiles and the metastatic potential of the tumors, as reported by earlier studies (21,41).

We expand these findings to show that the distinct transcriptional states correspond with increasing intrinsic degrees of resistance to MAPK/ERK pathway inhibition. Specifically, the melanocytic-like cell lines were prone to induced apoptosis upon targeting of BRAF and MEK, while elevated levels of resistance were found for the highly migratory mesenchymal-like state. More interestingly, our data provide new evidence indicating that tumors with an intermediate phenotype might already exhibit a considerable drug tolerance, despite not having yet fully developed migratory properties and showing mixed levels of expression of genes from invasive melanoma signatures. A significant population of quiescent cells, however, could already be detected in the intermediate state, congruent with previous findings linking a slow-cycling phenotype with residual disease and therapy resistance (42). Indeed, we could observe an overall good correspondence between the degree of quiescence and resistance in our panel of cell lines.

Our results additionally imply that the decrease in the sensitivity to drugs targeting the MAPK/ERK pathway is also delineated by a clear loss of transcriptional signatures of melanocytic differentiation, supporting previous findings (29). Further insights to this were revealed by information properties of the transcriptomes, being particularly interesting to note that the intermediate state cell line exhibited the highest transcriptome diversity. This is not only to be expected from tumors displaying mixed melanocytic and mesenchymal features, but it also suggests that the intermediate phenotype might represent a pluripotent cellular state that confers the cells with the plasticity required to subsequently evolve and trans-differentiate to adapt to drug exposure.

The implementation of single-cell approaches has allowed to recognize the intermediate phenotype as a distinct cellular state bearing unique features, rather than a mixed population of melanocytic- and mesenchymal-like cells (41). As we and others have observed, this state has the particularity of exhibiting a high expression of neural crest-like and immune-related transcriptional programs, including signatures of tumor necrosis alpha (TNF-α) and interferon gamma (IFN-γ) signaling (29,41). These pro-inflammatory pathways have been previously related to induction of dedifferentiation, slow-cycling phenotype, and diminished sensitivity to MAPK/ERK pathway inhibition in melanoma (29,43–45). Hence, these features might promote intermediate-state cells to further transition into more dedifferentiated phenotypes.

A reduction of melanocytic differentiation may therefore not only be relevant to regain properties from neural crest cells, which are characterized as highly migratory and multipotent (46), but may constitute an initial step required to undergo further transformation and successfully develop new malignant features. In line with this, and alike others (29,41), we observed that melanoma cells can undergo further dedifferentiation, losing neural crest features that are prominent in the intermediate state, such as expression of nerve growth factor receptor (NGFR), Erb-B2 receptor tyrosine kinase 3 (ERBB3), and SOX10. It has been shown that the transcriptional specialization of tumors is significantly less than for the corresponding normal tissues and comparable with that of dedifferentiated embryonic stem cells, which is partially attributed to a decrease in the expression of genes that are highly specific to normal cells (30). However, we show that cancer cells with very high resistance and metastatic potentials might display more specialized transcriptomes in comparison to other tumors in earlier stages of progression, reflecting the acquisition of novel malignant features. In line with this, we observed that highly resistant melanoma cells can even display greater transcriptome resemblances with fibroblasts than with non-transformed melanocytes or other more sensitive/differentiated melanoma tumors, hinting that trans-differentiation processes might give rise to highly resistant cell populations.

The large transcriptional reprogramming that tumor cells can undergo during late stages of malignant progression, as we observed, entails the activation of compensatory signaling cascades. This allows for the development of intrinsic resistance due to indifference to MAPK/ERK pathway inhibition, as opposed to incomplete inhibition. This might explain why the response to vemurafenib could not be improved by co-targeting these transporters, despite the most resistant cell line displaying stem-like properties with enhanced efflux capacity due to over-expression of multidrug resistance ABC transporters. Acquired resistance to kinase inhibitors has been previously shown to correlate with an increased expression of specific ABC transporters, such as ABCB1 and ABCG2, for which vemurafenib is known as a substrate (36). Our data, however, argue against ABC transporter-mediated efflux acting as a prominent mechanism of intrinsic resistance to MAPK/ERK targeted therapies, through reduction of intracellular concentrations of the drugs. This, together with lower basal levels of transcriptional signatures of MAPK/ERK pathway activity, suggests that a reduced dependency on this signaling cascade might be more relevant during late stages of resistance. These findings might partially account for the limited clinical relevance that pharmacological targeting of ABC transporters has proven (47,48).

Independency from MAPK/ERK signaling could be facilitated to some extent by shifting to a slow-cycling/dedifferentiated phenotype. The role of BRAF as driver of melanocytic transformation has been linked in great part to its function in transducing c-Kit signaling, which constitutes a main regulator of normal melanocyte survival and proliferation (49,50). Hence, during early stages of oncogenesis, activating mutations in BRAF or other MAPK/ERK pathway components might contribute to malignant transformation by inducing abnormal melanocytic proliferation. A phenotypic switch into a more-quiescent/highly-migratory phenotype, supported by the activation of alternative signaling cascades, might allow independency of this signaling pathway at the latest stages of progression, facilitating a better survival under therapeutic pressures.

Although several studies have converged upon reactivation of the MAPK pathway as a relevant mechanism of acquired resistance in melanoma (51–54), our findings indicate that increased compensatory signaling through the PI3K/AKT/mTOR axis might be generally more determinant in driving late-stage resistance. Indeed, combined targeting of this pathway has been shown to improve the response to MAPK/ERK inhibition (55) and lack of PTEN expression, a critical negative regulator of the PI3K activity that was absent in our prototypical highly-resistant/mesenchymal-like cell line, can be found in up to 30% of melanomas, often concurrent with the BRAF^*V600E*^ mutation (56).

Remarkably, we observed that PI3K/AKT/mTOR signaling sustained stem cell features from melanoma cells, as suggested by the abrogation of the SP by the dual PI3K/mTOR inhibitor dactolisib. In line with this, previous studies have also linked an increased PI3K/mTOR activity with dye exclusion abilities in breast cancer (57) and glioblastoma (58). In these studies, dye efflux was enhanced by lack of PTEN and was conversely reversed by the knockdown of mTOR as well as pharmacological PI3K or mTOR inhibition. In melanoma, pharmacological inhibition of PI3K has been shown to subdue TNF-driven sphere formation abilities from the cells, suggesting that PI3K/AKT signaling might mediate the inhibition of melanoma differentiation induced by pro-inflammatory pathways (59). Moreover, it has been demonstrated PI3K/AKT signaling is required for the TGF-β-induced EMT-like conversion of melanoma cells (60). Altogether, this suggests a clinical potential of targeting the PI3K/AKT pathway to overcome resistance driven by loss of differentiation.

## Conclusions

Although the current therapeutic management of patients with advanced melanoma is highly dependent on the mutational state of BRAF, a large phenotypic and molecular heterogeneity can be observed among tumor cells bearing the same BRAF mutation. This provides a multifactorial basis for intrinsic resistance, which corresponds with the loss of differentiation, increased quiescence and migratory potential, and the activation of alternative signaling pathways. In particular, the activation of the PI3K/AKT pathway in mesenchymal-like tumors confers independency from MAPK/ERK signaling, rendering inhibitors from the latter pathway largely ineffective. More personalized combined treatment approaches should be developed to target compensatory signaling pathways as a way to overcome late-stage resistance.

## Declarations

## Acknowledgements

The authors express their gratitude to Ms. Prakriti Shrestha for her technical collaboration, to Nikolai Lontke and Geritt Batt for their support with the computing infrastructure management, and to the high throughput sequencing unit team of the DKFZ Genomics and Proteomics Core Facility, Heidelberg, for providing the microarray services.

## Authors’ contributions

Conceptualization, MB, EC, EL-Z, PM; methodology, EC, EL-Z, PM; investigation, EC, EL-Z, RM, AL, SL, SK; formal analysis, EC, EL-Z, PM, JM; visualization, EC, EL-Z; writing, EC, EL-Z; review and editing, EC, EL-Z, MB; resources, MB; funding acquisition, MB. All authors have read and approved the final manuscript.

## Funding

This project was funded by the German Research Foundation (Deutsche Forschungsgemeinschaft, DFG) Research Training Group GRK2344 “MeInBio-BioInMe” (to EC, RM, and MB), SFB1479 (Project ID: 441891347-S1), SFB1160 (Project Z02), SFB1453 (Project ID 431984000 - S1) to MB, and the German Federal Ministry of Education and Research (Bundesministerium für Bildung und Forschung, BMBF) by MIRACUM within the Medical Informatics Funding Scheme (FKZ 01ZZ1801B to MB and PM).

## Availability of data and materials

The datasets generated during the current study will be publicly available by the time of publication. The following datasets were downloaded from the GenBank Sequence Read Archive: SRR3192558, SRR3192559, SRR3192555, SRR3192554, SRR1182320, SRR1182319, SRR1182317, SRR1182316 (HEM), SRR16890873, SRR16890874, SRR16890889, SRR16890890 (MaMel21), SRR16890875, SRR16890876, SRR16890891, SRR16890892 (MaMel63a), and GSE80829 (panel of 53 melanoma cell lines).

## Competing interests

The authors declare that they have no competing interests.

## Notes

### Competing Interest Statement

The authors have declared no competing interest.

