## Supplementary figures for "PI3K/AKT signaling allows for MAPK/ERK pathway independency mediating dedifferentiation-driven treatment resistance in melanoma"

#### Fig. S1

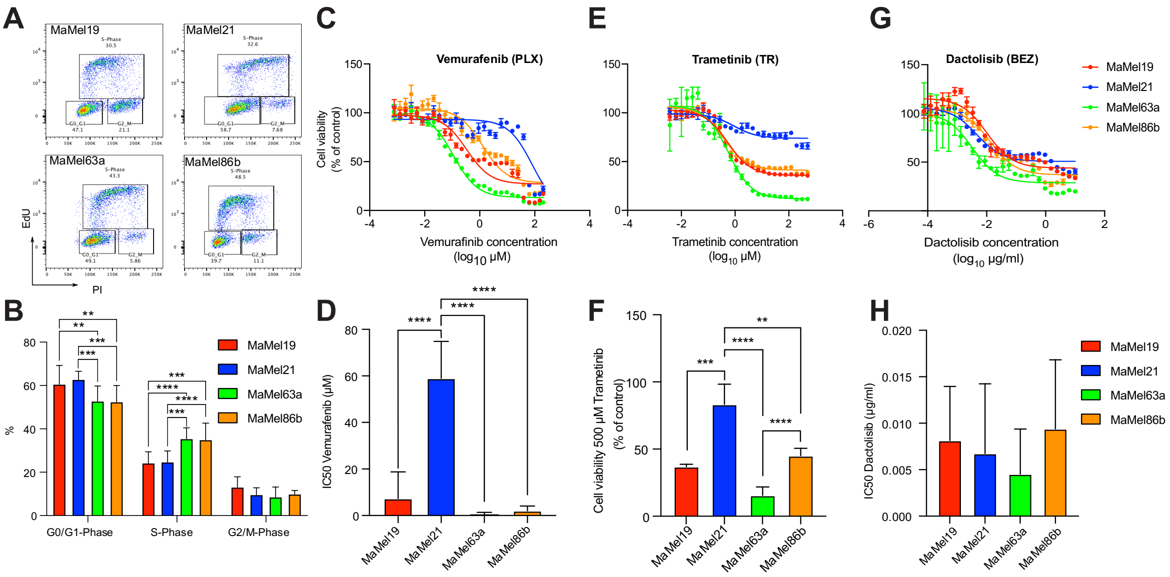


**Fig. S1. Cell cycle analysis and sensitivity to inhibitors in BRAF$^{V600E}$ mutant MaMel cells.**

(**A**) Representative flow cytometric analysis of EdU and PI intracellular co-staining, defining the cell cycle phases G0/G1, S, and G2. (**B**) Quantification of frequencies of the cells in cell cycle phases shown in (A). Data from five independent experiments is shown (n = 5). **(C)** Representative analysis of MaMel cell viability under inhibition with vemurafenib (PLX; concentration range: 200 µM - 0.8 nM). (**D**) IC50 values obtained from MTT assays on MaMel cells under inhibition with PLX. Data from five independent experiments are shown (n = 5). (**E**) Representative analysis of MaMel cell viability under inhibition with trametinib (TR; concentration range: 1 µM - 0.002 nM). (**F**) Quantification of MaMel cell viability under 500 µM of trametinib. Data from three independent experiments are shown (n = 3). **(G)** Representative analysis of MaMel cell viability under inhibition with dactolisib (BEZ; concentration range: 10 µg/ml - 0.07 ng/ml). Data shown in C, E, and G were acquired using MTT assays. All bar plots show the mean ± SD of the individual experiments. *p < 0.05, **p < 0.01, ***p < 0.001, ****p < 0.0001.

### Fig. S2

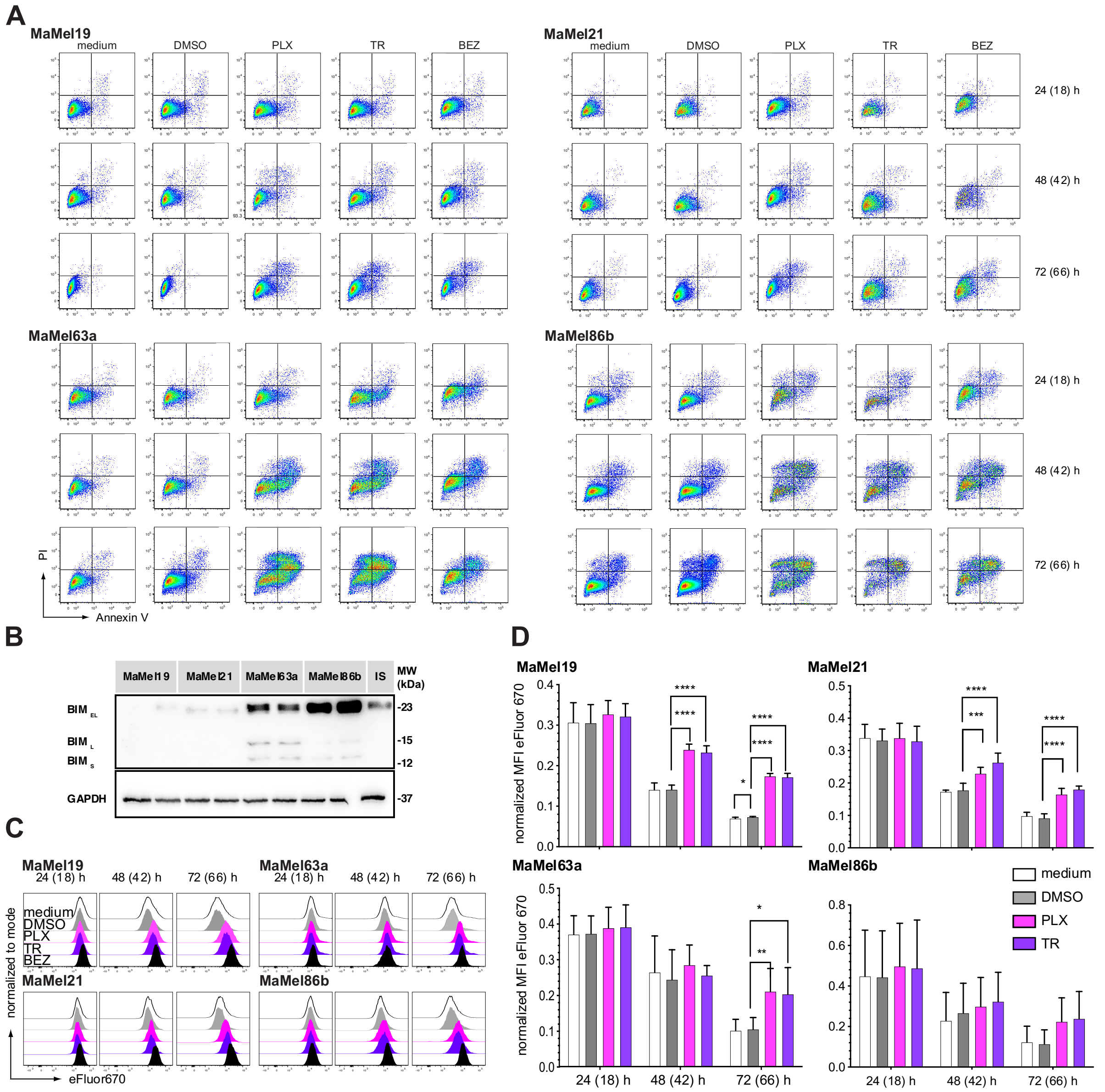


**Fig. S2. Apoptotic and proliferative response of BRAF$^{V600E}$ mutant MaMel cells under inhibition.**

**(A)** Representative flow cytometric analysis of Annexin V and PI co-staining, defining the viable (Annexin V – PI negative; lower left quadrant) and apoptotic/necrotic cells in MaMel cell lines under the indicated treatment conditions for three different time points. BEZ = dactolisib, PLX = vemurafenib, TR = trametinib. (**B**) Western blot analysis of BIM expression in MaMel cell lines. Three BIM isoforms (EL, L, and S) are resolved. GAPDH served as loading control. (**C**) Representative flow cytometric analysis of cell proliferation rate using the eFluor 670 dye in MaMel cells under the indicated treatment conditions for three different time points. Cell proliferation is detected as a reduction of the fluorescence intensity of eFluor 670 dye. In (A) and (C) the time points outside the brackets indicate the total time of cells in culture. Time points in brackets indicate the inhibition time. (**D**) Quantification of the cell proliferation rate of MaMel cells under inhibition with PLX (3 µM) and TR (5 nM). Median fluorescence intensity (MFI) of the eFluor 670 dye, measured by flow cytometry, is depicted at the indicated time points. Proliferation is detected as a reduction of MFI. Data from three independent experiments are shown (n = 3). All bar plots show the mean ± SD of the individual experiments. $*p < 0.05$, $**p < 0.01$, $***p < 0.001$, $****p < 0.0001$.

#### Fig. S3

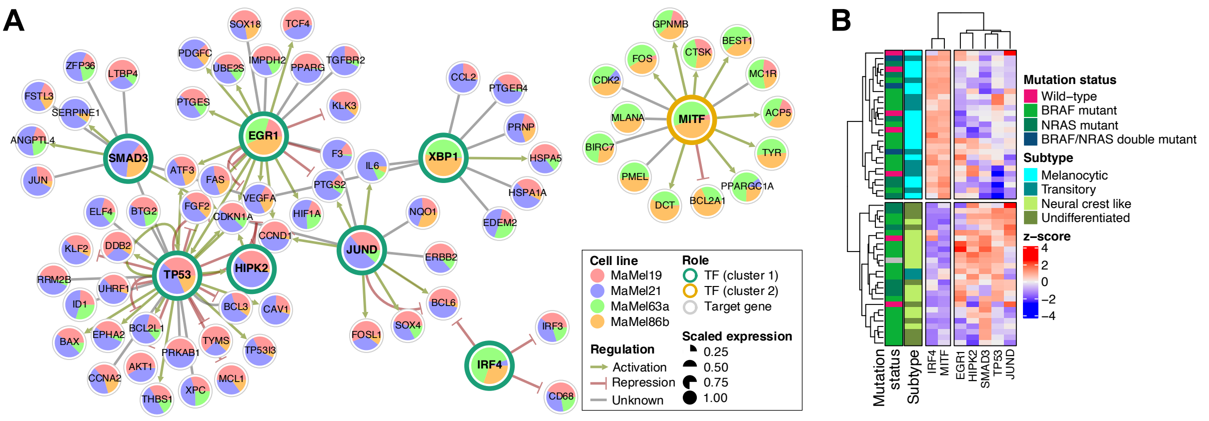


**Fig. S3. Over-represented targets of transcription factors within groups of genes correlated with the degree of resistance in BRAF*V600E* mutant MaMel cells.**

(**A**) Network of transcription factors (TF) and their target genes, enriched within clusters of genes increasing (cluster 1) or decreasing (cluster 2) their expression along with higher levels of resistance to MAPK/ERK pathway inhibition. The nodes are colored according to the scaled expression of the respective gene for the different MaMel cell lines. The arrow type and color of the edges indicate the predicted regulation exerted by the TF over the corresponding target gene, according to TRRUST v2 database (13). (**B**) Normalized expression (z-score) from TF in 53 melanoma cell lines from Tsoi et al. (29), showing different NRAS/BRAF mutation status and degrees of differentiation.

### Fig. S4

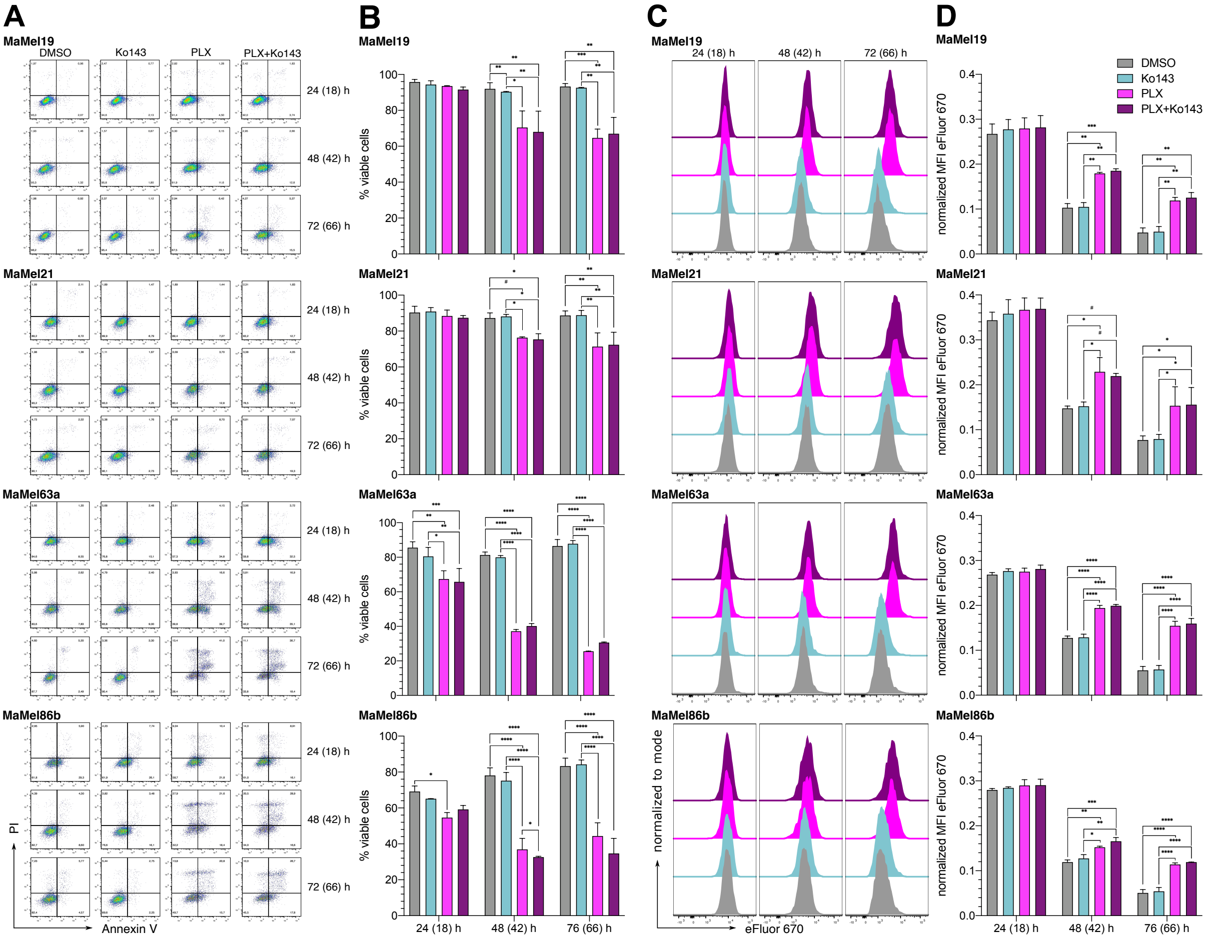


**Fig. S4. Apoptotic and proliferative response of MaMel cells under combined BRAF and ABC transporter inhibition.**

(**A**) Flow cytometric analysis of the apoptotic response of MaMel cells under inhibition with vemurafenib (PLX), ABCG2 inhibitor Ko143, or their combination (PLX+Ko143) for three different time points. (**B**) Representative Annexin V and PI co-staining, defining the viable (Annexin V – PI negative; lower left quadrant) and apoptotic/necrotic cells under the indicated treatment conditions. Right: quantification of the percentage of viable cells. (**C**) Flow cytometric analysis of cell proliferation rate of MaMel cells under inhibition with PLX, Ko143, or their combination for three different time points. Proliferation is detected as a reduction of the fluorescence intensity of eFluor 670 dye. (**D**) Representative histograms showing the dye fluorescence under the indicated treatment conditions. Right: quantification of median fluorescence intensity (MFI) of the eFluor 670 dye. Time points outside the brackets indicate the total time of cells in culture; time points in brackets indicate the inhibition time. All bar plots show the mean ± SD of two independent experiments (n = 2). $^{\#}p < 0.08$, $^{*}p < 0.05$, $^{**}p < 0.01$, $^{***}p < 0.001$, $^{****}p < 0.0001$.

### Fig. S5

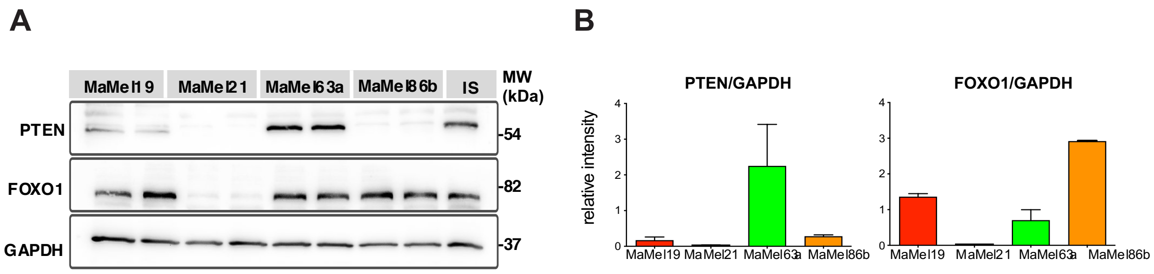


**Fig. S5. PTEN and FOXO1 expression in BRAF$^{V600E}$ mutant MaMel cells.**

(**A**) Representative Western blot analysis of PTEN and FOXO1 expression in MaMel cell lines. GAPDH served as a loading control. (**B**) Densitometry quantification for two independent experiments (n = 2). Optical density was normalized to the corresponding loading control and internal standard (IS). Bar plots show the mean ± SD of the experiments.
